# A copper chaperone-mimetic polytherapy for SOD1-associated amyotrophic lateral sclerosis

**DOI:** 10.1101/2021.02.22.432389

**Authors:** McAlary L., Shephard V.K., Wright G.S.A., Yerbury J.J.

## Abstract

Amyotrophic lateral sclerosis (ALS)-associated mutations in Cu/Zn superoxide dismutase (SOD1) reduce folding stability, resulting in misfolding, aggregation, and ultimately cellular toxicity. A great deal of effort has focused on preventing the misfolding and aggregation of SOD1 as a potential therapy for ALS, however, the results have been mixed. Here, we utilise a small-molecule polytherapy of CuATSM and ebselen to mimic the metal delivery and disulfide bond promoting activity of SOD1’s cellular chaperone, the ‘copper chaperone for SOD1’ (CCS). We find that polytherapy using CuATSM and ebselen is highly effective at reducing inclusion formation in a cell model of SOD1 aggregation, reduces mutant SOD1-associated cell death, and promotes effective maturation of SOD1 beyond either compound alone. Our data suggest that a polytherapy of CuATSM and ebselen may be an effective method of treating SOD1-associated ALS.

## Introduction

Over 160 mutations have been identified throughout the gene encoding Cu/Zn superoxide dismutase (SOD1) that are known to cause the motor neuron disease amyotrophic lateral sclerosis (ALS) [1,2]. These mutations result predominantly in amino acid substitutions found in all SOD1 protein secondary structure elements, cofactor-binding sites and homodimer interface. Each is thought to cause ALS by decreasing SOD1 folding stability, thereby creating a pool of misfolded and aggregation-prone SOD1 [3,4]. While there is debate as to whether proteinaceous aggregates or smaller, soluble non-native oligomers are toxic [5–7] the initial event considered to spark cell death is SOD1 protein misfolding [3,4,8–11].

SOD1 maturation comprises several sequential post-translational modifications (PTMs). Initially, the spontaneous binding of zinc (Zn) to immature monomer provides some folding stability [12]. Zinc-bound SOD1 then associates with the copper chaperone for SOD1 (CCS) which facilitates the input of copper (Cu) and subsequent oxidation of an intra-subunit disulfide bond between Cys57 and Cys146. The stable monomer is then free to form enzymatically active homodimers [13–15]. Metal-binding region (MBR) mutants affect copper or zinc coordination and activity while wild-type-like (WTL) mutants retain high levels of enzymatic activity when mature [16,17]. Immature SOD1, lacking PTMs, is prone to misfolding and is the central component of intracellular aggregates found with ALS neuronal tissues [3,8,10,11,18–20], whereas mature SOD1 is highly stable [21]. Maturation and misfolding are therefore antagonistic pathways that dictate SOD1 toxicity. A cell has finite resources and a limited capacity to catalyze nascent SOD1 maturation. The maturation pathway can be overwhelmed by high concentration nascent SOD1 [22] or inhibited by mutations that prevent cofactor binding and disulfide PTMs [3,4,9,23,24]. This results in increased misfolding pathway flux and the proteostasis pathways becoming overwhelmed [25]. Decreasing SOD1 expression and thereby reducing traffic along the misfolding pathway is the focus of knock-down strategies currently in clinical trials [26]. However, SOD1 has important metabolic functions and long-term ablation of its activity is known to have detrimental effects. A second option is to increase maturation or cellular proteostasis capacity. Heat shock protein molecular chaperone upregulation reduces SOD1 misfolding and clinical trials are again ongoing [27]. In addition, several small molecules have been shown to act as direct pharmacological chaperones for SOD1 [28–31]. Pyrimidine derivatives, 5-fluorouridine and telbivudine, reduce SOD1 *in vitro* aggregation and *in vivo* toxicity respectively [28,29]. Treatment of SOD1-G93A mice with the 5-fluorouridine analogue, 5-fluorouracil, also delays symptom onset and increases survival[32].

The cognate chaperone of SOD1, CCS, is a uniquely placed chaperone that has evolved to increase SOD1 maturation pathway throughput. It exerts molecular, copper, and oxidative folding chaperone activity on nascent SOD1 mediated through a specific protein-protein interaction [13,14,33,34]. While overexpression of hCCS reduces the accumulation of misfolded SOD1 it can also recruit SOD1 mutants to the mitochondrial intermembrane space where they accelerate vacuolization and toxicity [35–37]. Two small molecules have been shown to recapitulate CCS activities. Copper(II)ATSM (CuATSM) promotes WTL mutant SOD1 Cu-binding in several mouse models and increases lifespan [31,38–40] but it is ineffective against MBR mutants [41]. The seleno-organic compound ebselen promotes the formation of the SOD1 intra-subunit disulfide bond in cultured cells [30] and increases SOD1 dimer affinity [42,43]. As there are few SOD1 ALS-associated mutations that directly prevent disulfide formation (C146R and truncation mutants) ebselen is likely to be effective for MBR mutants as well as WTL [2]. Indeed, recent evidence shows that ebselen and some of its derivatives can restore the viability of cultured cells expressing G93A mutant SOD1, as well as delay disease onset in G93A mice through dietary supplementation [44].

Here, we report the effect of ebselen on intracellular mutant SOD1 inclusion formation in a disease-relevant cell model. To achieve this, we developed a machine learning-based image analysis pipeline for accurate measurement of protein inclusion formation in large microscopy data sets. Application of this method to a subset of compounds showed ebselen was capable of reducing inclusion formation for both WTL and MBR SOD1 mutants. We then utilized this method to investigate CuATSM and ebselen co-therapy aiming to divert nascent SOD1 from the misfolding pathway to the maturation pathway and thereby reduce mutant toxicity. We show these compounds can act in a synergistic manner to reduce SOD1 aggregation through the promotion of dimerization, disulfide formation and copper loading. All mutants analyzed display positive outcomes for at least one marker of effective SOD1 maturation with resulting cell viability increases for common or severely structurally destabilizing mutants. This work highlights the unexplored possibilities of mutation-specific personalized therapy for SOD1-ALS and the potential use of a CCS-mimetic polytherapy specifically targeting steps on the SOD1 PTM maturation pathway.

## Results

### Automated image analysis to identify cells containing inclusions

We, and others, have previously utilized genetically encoded fluorescent proteins as a tool to investigate the inclusion formation of SOD1 in cultured cells finding that ALS-associated and *de novo* mutations, as well as small molecules, can alter this process [3,29,41,45–48]. We have also utilized several methods to detect fluorescent proteinaceous inclusions, including manual counting [3], fluorescence intensity thresholding [29], and cell permeabilization to release soluble GFP-tagged protein [29,41,49]. We now sought to enhance our detection of protein inclusion formation in cells for use in larger mutational or drug screens. This was accomplished by exploiting advances made in the area of microscopy image analysis with the application of user-assisted machine learning to accurately classify cellular phenotypes [50].

To this end, we developed an image analysis pipeline using CellProfiler software [51] to identify and segment transfected cells for measurement followed by classification in CellProfiler Analyst software. Measurement parameters were chosen to append spatial data (shapes, texture, granularity, radial intensity, and intensity) in order to generate cytoprofiles that were processed by a random forest classifier **(Figure 1A and Supp. Fig. 1 and 2A)** [50]. Normalization of the extracted cellular features demonstrated significant differences in texture, radial intensity, and intensity measurements, but not shape or granularity **(Supp. Fig. 2B)**. This indicated that these measurements were appropriate for the profiling of cells. We found that our segmentation parameters correctly identified transfected cells with high accuracy (97 ± 2%) and that the chosen measurements effectively facilitated accurate classification for both cells with and without inclusions (95 ± 0.5% and 98 ± 0.9% accuracy respectively) **(Figure 1B)**.

**Figure 1.**
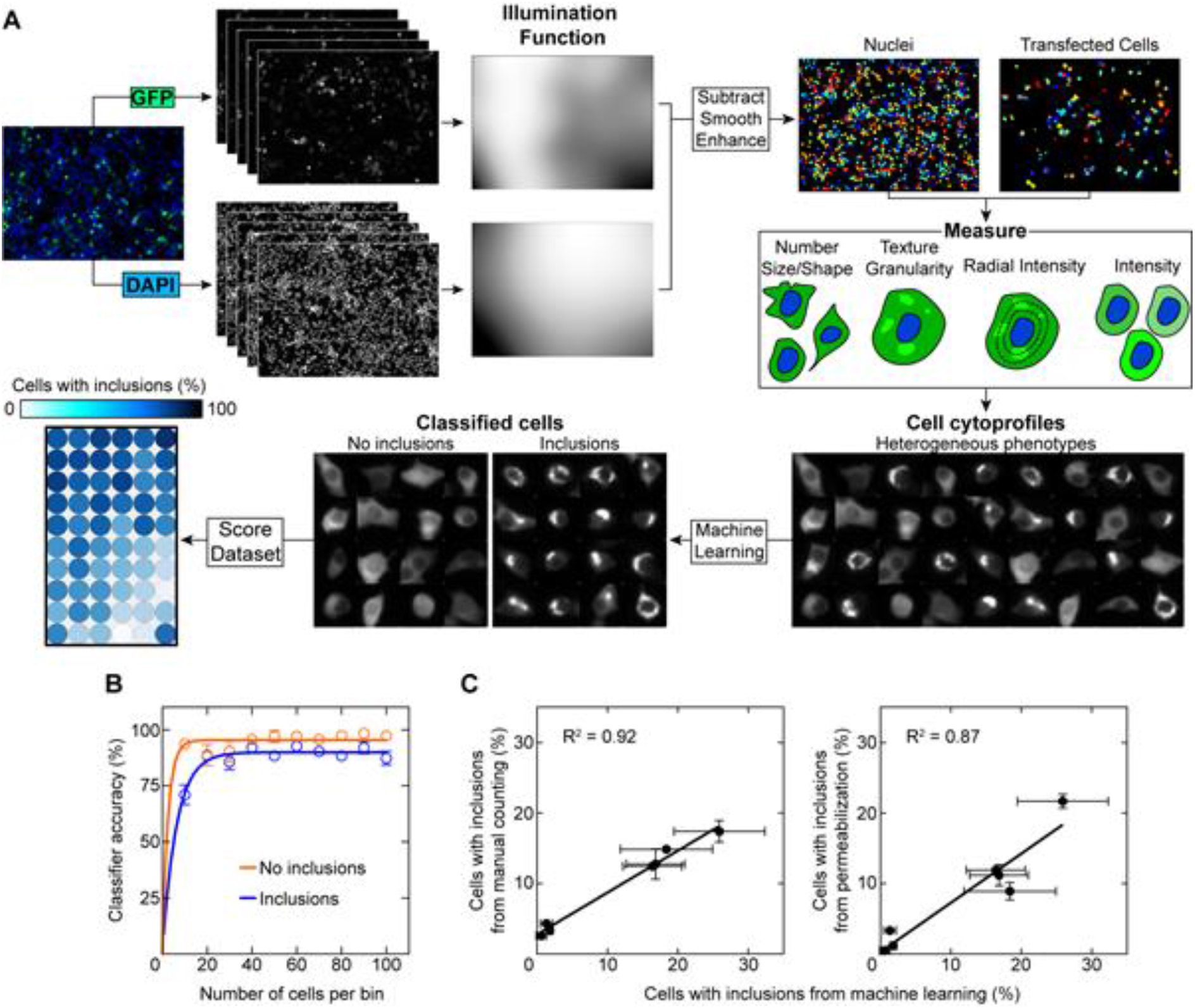
User-assisted machine learning to determine cells containing inclusions. **(A)** The image analysis pipeline first performs illumination correction for both DAPI and GFP channels and then segments the transfected cells for measurement. A user then identifies phenotypes in a small subset of the cell population to train the classifier for identification of the entire cell population. **(B)** The classifier requires less than 100 to become accurate at categorizing cells into inclusion containing (blue) and those that did not contain inclusions (orange). **(C)** Correlations of the percentage of cells with inclusions in this work *vs* previously published examination of the same SOD1-EGFP expression constructs in NSC-34s. **(Left)** *vs* manual counts from McAlary et al. 2016 [3], and **(right)** *vs* saponin permeabilized cells from Farrawell et al. 2018 [25]. Error bars represent SD of the mean from at least 3 separate classification requests.

**Figure 2.**
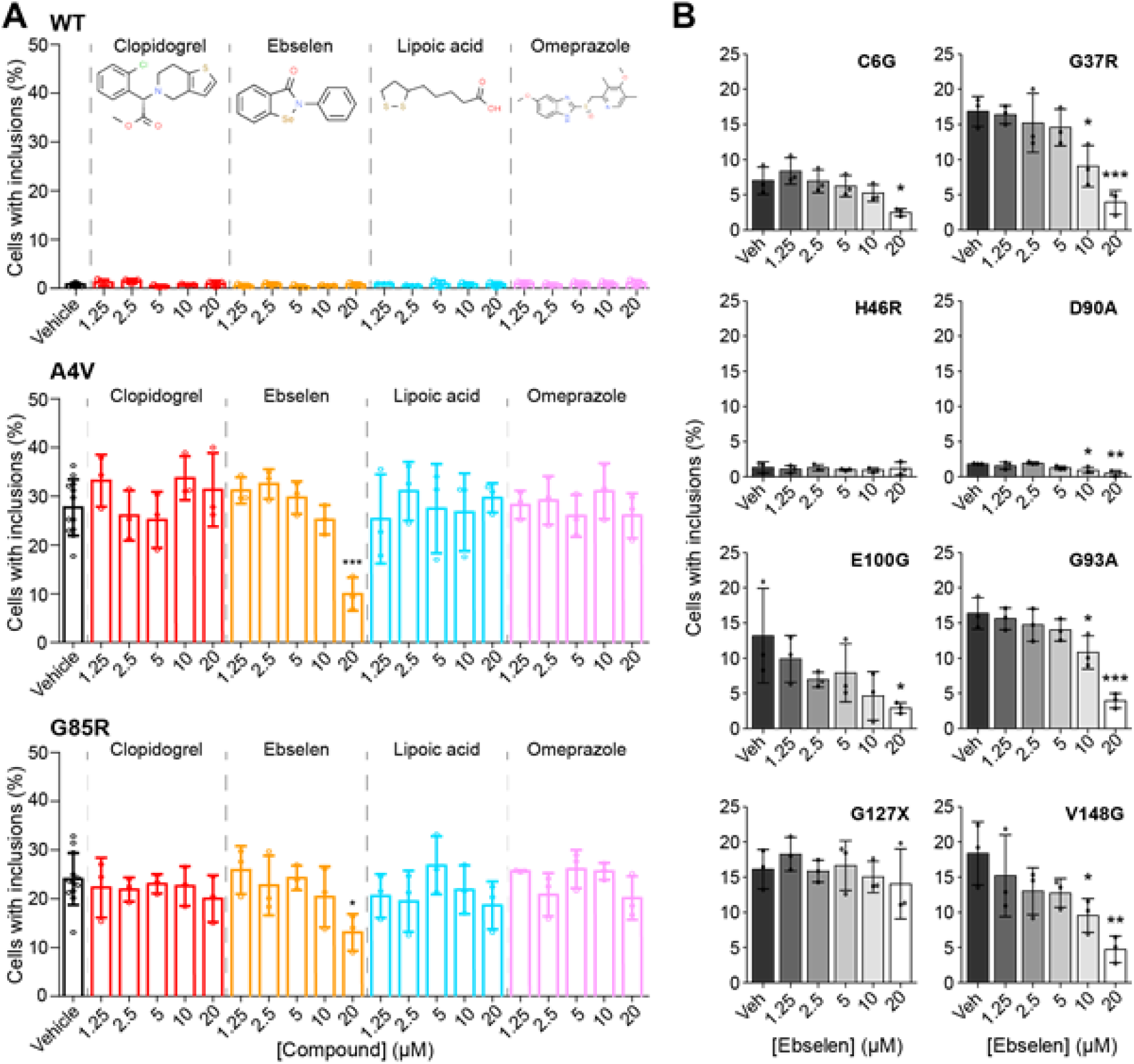
Ebselen reduces inclusion formation of SOD1 ALS-associated mutants in cultured cells. NSC-34 cells expressing **(A)** SOD1-EGFP variants WT, A4V, and G85R were treated with vehicle (black), clopidogrel (red), ebselen (orange), lipoic acid (teal), or omeprazole (pink) for 48 h and the number of cells containing inclusions was enumerated. **(B)** NSC-34 cells expressing SOD1-EGFP variants C6G, G37R, H46R, D90A, E100G, G93A, G127X, and V148G were all treated with increasing concentrations of ebselen and inclusion formation was measured. Ebselen treatment decreased inclusion formation for most variants except for G127X and H46R. Error bars represent SD of the mean of at least 3 separate experiments. Statistical significance was determined using a student’s t-test against vehicle control (*p < 0.001* = ***, *p < 0.05* = *).

### Ebselen, but not other compounds, reduces the formation of ALS-associated mutant SOD1 inclusions in cultured cells

Having established a rapid and accurate image-based method of classifying cells containing inclusions, we next sought to examine the effect of ebselen and a small panel of other similar small molecules on SOD1 inclusion formation. The molecules, other than ebselen, were omeprazole, clopidogrel, and lipoic acid. These compounds were chosen on the basis that they contained sulfur moieties that may be redox-active in a similar manner to ebselen [30].

To this end, NSC-34 cells were transfected with SOD1 variants WT, A4V (WTL mutant), or G85R (MBR mutant) and were treated with determined non-toxic concentrations of ebselen, lipoic acid, omeprazole, or clopidogrel **(Supp. Fig. 3)** for 48 h prior to being fixed, imaged, and analysed. We observed that WT SOD1 formed very few inclusions across treatments, with only 0.8 ± 0.3% of cells in the untreated group being classified as containing inclusions **(Figure 2A)**, in line with previous observations by us and others [3,29,41,45–48]. In contrast, both A4V and G85R SOD1 readily formed inclusions in this system with 27.7 ± 5.8% and 24.1 ± 5.3% cells containing inclusions respectively **(Figure 2A)**. Treatment of A4V or G85R SOD1 expressing cells with either lipoic acid, omeprazole, or clopidogrel at any of the tested concentrations had no significant effect on inclusion formation **(Figure 2A)**. We found that treatment with ebselen at the highest concentration of 20 μM resulted in a significant reduction in both A4V and G85R inclusion formation **(Figure 2A)**, suggesting that ebselen was protective against both WTL and MBR mutant inclusion formation in this model. The reduction in A4V mutant inclusion formation was more substantial (3-fold decrease) when compared to G85R (2-fold decrease). Previous examination of the capability of ebselen to facilitate SOD1 maturation in cells used a ten-fold greater concentration of ebselen (200 μM) than we used here [30], where our results suggest that ebselen may be more potent at facilitating SOD1 maturation than previously suggested.

**Figure 3.**
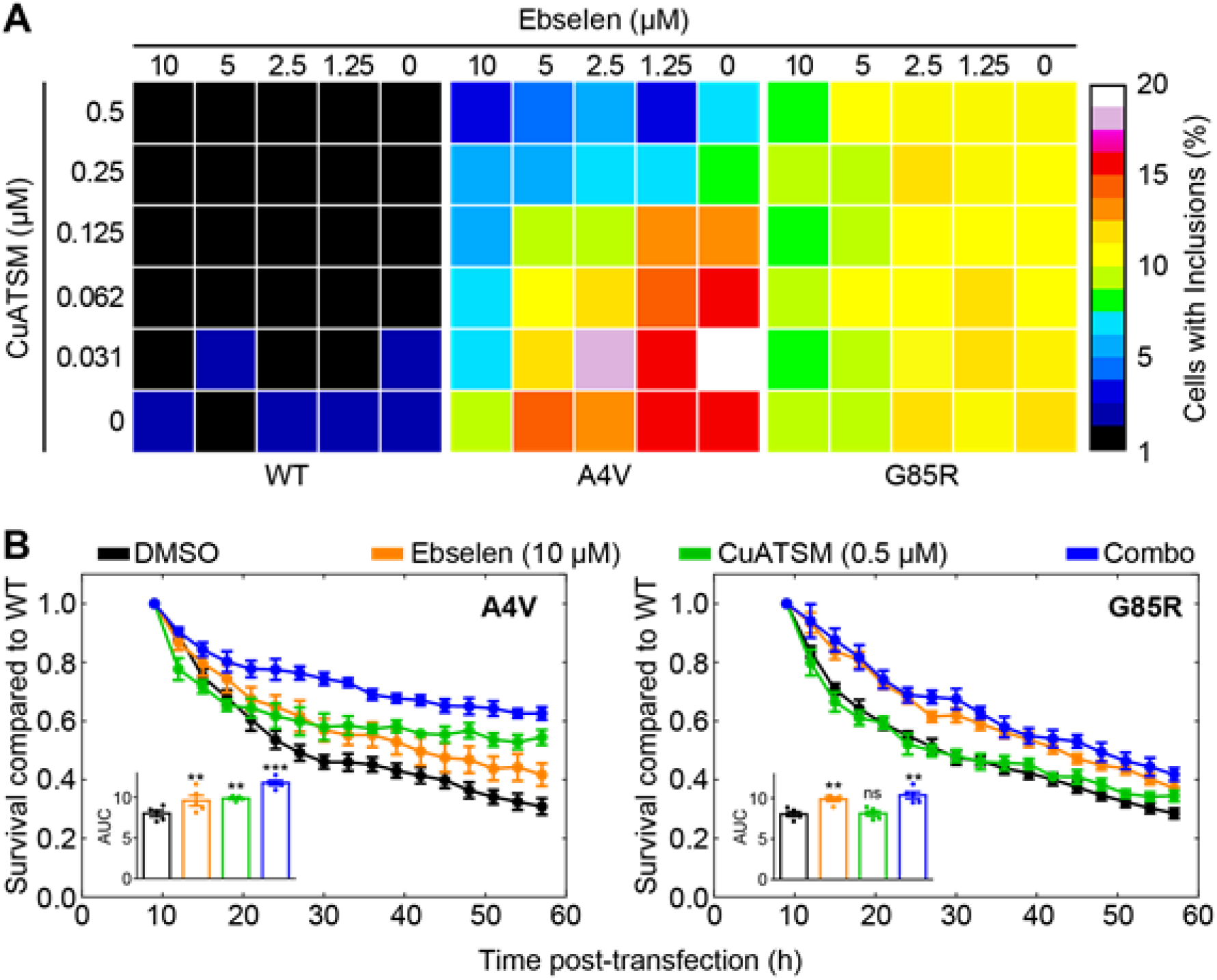
A combination treatment of CuATSM and ebselen is effective at rescuing SOD1-A4V folding. **(A)** Heatmaps of CuATSM and ebselen checkerboard treatment of NSC-34 cells expressing either SOD1-WT (left), SOD1-A4V (middle), and SOD1-G85R (right). Colours represent the mean percentage of transfected cells containing inclusions from 3 separate experiments. **(B)** Transfected cell counts of NSC-34 cells expressing SOD1-A4V (left) and SOD1-G85R (right) treated with vehicle DMSO (black), ebselen (10 μ M; orange), CuATSM (0.5 μ M; green), and ebselen/CuATSM combo (10 μ M/0.5 μ M; blue). Cell counts are relative to SOD1-WT transfected cells treated with the same compounds. Inset in each panel is the area under the curve measurements for each drug treatment. Error bars represent SEM of 3 separate experiments. Statistical significance was determined using a one-way ANOVA with Dunnet’s test against DMSO-treated cells (*p < 0.01* = ***, *p < 0.05* = **).

Ebselen was further tested on SOD1 variants including C6G, G37R, H46R, D90A, G93A, E100G, G127X, and V148G in this model. Most of these ALS-associated mutants in this list are WTL and induce inclusion formation to various degrees in NSC-34 cells [3,41]. H46R and G85R are MBR mutants with minimal ability to bind copper [52,53], and G127X is a truncation mutant that removes residues 127-153, including the disulfide forming Cys146 residue [54]. Treatment of the transfected NSC-34 cells with increasing concentrations of ebselen showed that for cells expressing C6G, G37R, D90A, G93A, E100G, and V148G there was a significant dose-dependent response to ebselen **(Figure 2B)**. The most effective dose observed in each case was 20 μM, although a significant difference between vehicle control and a concentration of 10 μM was observed for G37R, D90A, G93A, and V148G **(Figure 2B)**, indicating a greater effective action of ebselen on these mutants. Cells expressing the truncated G127X SOD1 mutant, which does not contain the disulfide-forming Cys146 residue, showed no significant effect of ebselen on the percentage of cells containing inclusions at any concentration tested **(Figure 2B)**. H46R transfected cells showed no response to ebselen, however, inclusion formation is low for this mutant. These data indicate that ebselen is likely acting through the previously proposed method of promoting the formation of the SOD1 intrasubunit disulfide between residues Cys57 and Cys146 [30].

### A Combination of Ebselen and CuATSM is Effective at Decreasing SOD1 Inclusion Formation in Cells

Multi-drug polytherapies are used as a standard treatment against most types of cancer [55]. This is a less common but growing strategy being adopted against neurodegenerative diseases [56,57]. Previous research has shown that the copper carrying compound CuATSM is capable of facilitating copper delivery to SOD1 in animals and cells [31,41]. Likewise, we have shown that ebselen can promote SOD1 disulfide formation [30]. Therefore, we reason that CuATSM and ebselen have the potential to act collaboratively to promote both copper binding and disulfide formation respectively in SOD1 mutants [30,31,41].

Considering this, we set out to establish if a combination of both CuATSM and ebselen would have a greater effect at reducing SOD1-associated fALS phenotypes in our cell model than either drug alone. To this end, we performed a checkerboard treatment of NSC-34 cells expressing WT, A4V, or G85R using different concentrations of CuATSM (0.5 - 0 μM) with ebselen (10 - 0 μM). Similar to our previous measurements with automated image analysis **(Figures 3A)**, WT formed very few inclusions in this assay at all CuATSM-ebselen combination treatments **(Figure 3A)**. In comparison, both A4V and G85R formed more inclusions at lower drug concentrations **(Figure 3A)**. Heatmap visualisation of the checkerboard analysis shows that A4V responds to both CuATSM and ebselen monotherapy, whereas G85R shows no response to CuATSM monotherapy but responds to higher concentrations of ebselen monotherapy. Interestingly, application of fractional inhibitory concentration index (FIC) measurement to the checkerboard assays showed that ebselen and CuATSM acted synergistically for A4V but not G85R. Indeed, ebselen at a concentration of 10 μM was capable of significantly reducing the number of cells with inclusions to roughly 5% with as little as 0.03 μM CuATSM, which is over a 10-fold decrease in the maximum CuATSM concentration. Likewise, a concentration of 0.25 μM CuATSM was capable of reducing the necessary ebselen concentration for significant inclusion formation reduction by roughly 8-fold, from 10 μM to 1.25 μM.

We next performed a live-cell time-lapse microscopy assay to count the relative numbers of GFP-positive cells across time under the most potent treatment regimes used in the checkerboard assay (CuATSM at 0.5 μM and ebselen at 10 μM). Data are reported as the number of cells that are EGFP-positive relative to SOD-WT-EGFP transfected cells treated with CuATSM, or ebselen, or a combination of both compounds. The expectation is that the compounds would reduce the time-dependent decline in relative GFP-positive cell numbers. Similar to previous reports [3,29,41,58], transfection of NSC-34 cells with mutant SOD1-GFP results in a decline in relative EGFP positive cells over time **(Figure 3B; left A4V, right G85R)**. In this assay, cells transfected with A4V and treated with either ebselen or CuATSM alone, or with the combination therapy saw a significant increase in the number of GFP-positive cells across time as determined by measuring the area under the curve **(Figure 3B; left and inset)**. Cells transfected with G85R also showed a similar trend of decreasing numbers of relative GFP-positive cells across time, however, CuATSM treatment alone had no effect on this decline whereas ebselen alone and ebselen with CuATSM in combination did significantly reduce the decline in cell numbers relative to WT **(Figure 3B; right and inset)**.

### Polytherapy with Ebselen and CuATSM mimics CCS Activity and is Effective at Promoting Mutant SOD1 Maturation

To investigate the mechanisms by which ebselen and CuATSM catalyse SOD1 maturation we assessed the intra-subunit disulfide bond formation, dimerization, and activity of SOD1. In comparison to the other redox compounds examined previously, only ebselen was able to facilitate purified recombinant A4V disulfide formation **(Figure 4A)**. Intra-subunit disulfide formation is known to shift the SOD1 monomer-dimer equilibrium in favour of the dimer [59] and we have previously shown ebselen binding to Cys111 can increase A4V homodimer affinity. However, this effect is negated by the presence of dithiothreitol or reduced glutathione at physiological concentrations of 1 mM [30]. Ebselen, unlike oxidized glutathione, was able to facilitate the formation of SOD1 homodimers even in the presence of 5 mM reduced glutathione **(Figure 4B)**. Under these conditions, ebselen cannot form stable conjugates at Cys111, therefore, SOD1 homodimerization likely results from the catalyzed formation of the SOD1 intra-subunit disulfide formation as we previously described in live cells [30].

**Figure 4.**
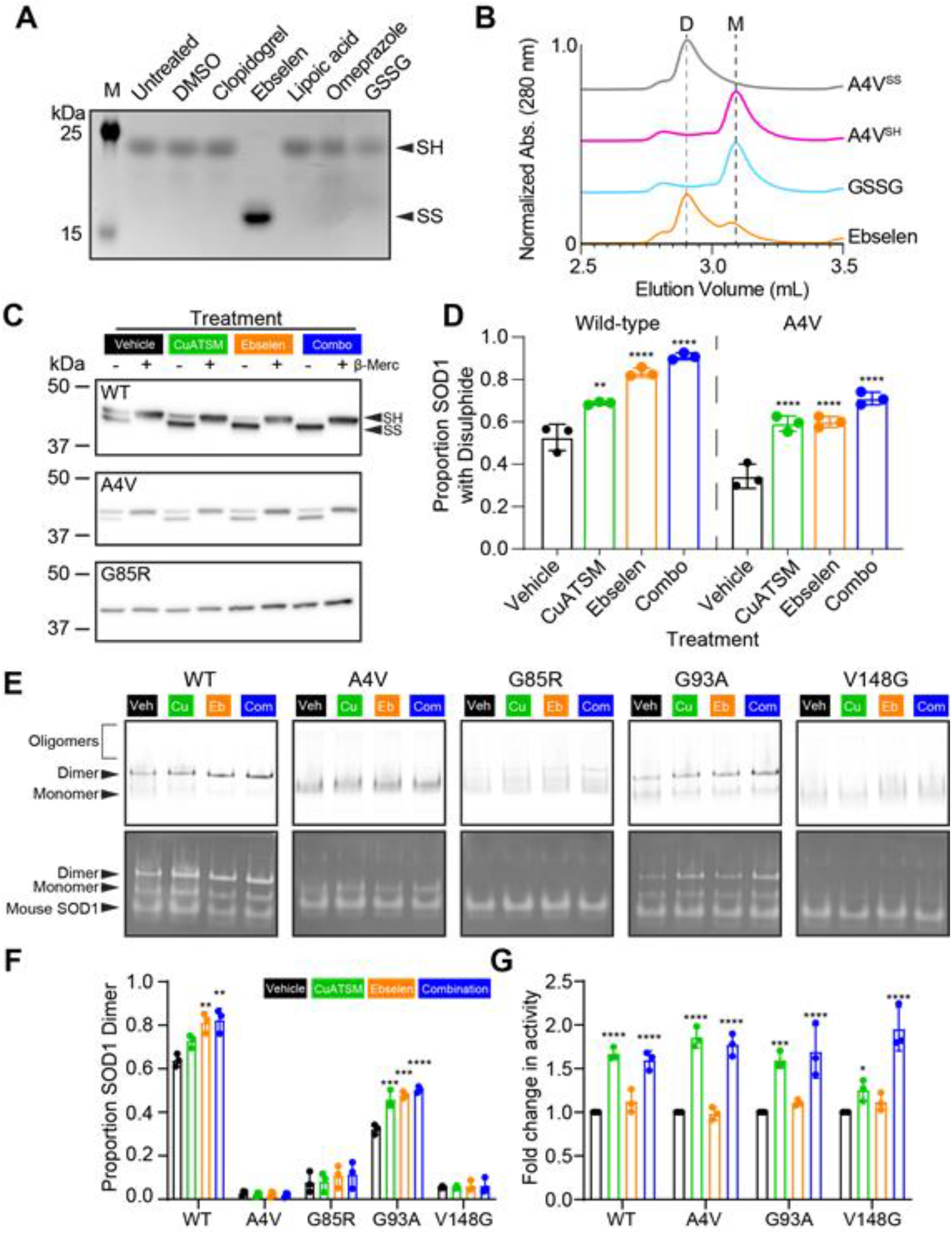
A combination treatment of CuATSM and ebselen is effective at rescuing SOD1 ALS-associated mutant folding. **(A)** Non-reducing SDS-PAGE AMS assay shows treatment with ebselen but not other redox containing compounds resulted in SOD1-A4V disulfide formation (reduced = SH, oxidized intact =SS). **(B)** Size exclusion chromatography shows ebselen promotes A4V homodimerization whereas oxidized glutathione (GSSG) does not (D = dimers, M = monomers). **(C)** Differential SDS-PAGE migration of SOD1-EGFP from cell lysates under reducing (+β-merc) and non-reducing (-β-merc) conditions shows that the proportion of disulfide bonded SOD1 (SS) is increased with CuATSM (green; 0.5 μ M), ebselen (orange; 20 μ M), and a combination treatment (blue; 0.5 μ M CuATSM/20 μ M ebselen) compared to vehicle control (black) for both WT and A4V, but G85R remains fully reduced. **(D)** Densitometry of disulfide formation immunoblots for WT and A4V showed that CuATSM, ebselen, and the combination therapy were capable of promoting disulfide formation in living cells. **(E)** Native-PAGE of SOD1-TdTomato lysates shows oligomers, dimers, and monomers of SOD1 variants (top) and in-gel zymography of the same gels shows the relative activity of each species including dimer, monomer, and mouse SOD1. **(F)** Quantification of the fluorescence signal from native-PAGE of the proportion of SOD1-TdTomato signal present for the dimer, showing that CuATSM, ebselen, or combination therapy promoted the dimerization of both WT and G93A SOD1, but not A4V, G85R, and V148G. **(G)** Quantification of the achromatic bands from in-gel zymography showing that only treatment with CuATSM and the combination therapy increased the relative levels of active SOD1 for WT, A4V, G93A, and V148G (G85R not shown due to lack of activity). Error bars represent SD of the mean of at least 3 separate experiments. Significance was determined using one-way ANOVA with Dunnet’s multiple comparisons test with comparisons made against vehicle control data (*p < 0.0001* = ****, *p < 0.001* = ***, *p < 0.01* = **, *p < 0.05* = *).

Next, we compared the effectiveness of CuATSM and ebselen monotherapies and combination therapy on their ability to promote the folding of intracellular SOD1 variants. Previous work determined that non-reducing SDS-PAGE of SOD1 maintains the intramolecular disulfide bond and that the disulfide bonded form of monomeric SOD1 migrates more rapidly during electrophoresis [14]. SOD1 containing the intra-subunit disulfide was detected across treatment groups for both WT and A4V, but not for G85R **(Figure 4C)**. Semi-quantitative measurement of the immunoblots showed a significant shift in the proportion of disulfide-containing SOD1 detected for both WT and A4V for each treatment in comparison to vehicle control **(Figure 4D)**. Out of the treatments, the combination treatment showed the greatest shift in the proportion of SOD1 containing the intra-subunit disulfide for both WT and A4V **(Figure 4C and D)**. The lack of detection of disulfide bonded G85R may be a result of this mutant being highly destabilized even in cells, and being highly susceptible to reduction even when free-thiols are chemically blocked [60–62].

We next examined the effect of CuATSM and ebselen monotherapies and polytherapy on SOD1 dimerization and activity in cells by using Native-PAGE and in-gel zymography [63]. Here we used SOD1 variants tagged with TdTomato due to EGFP-tagged SOD1 migrating too close to endogenous cellular SOD1 for accurate densitometry measurements **(Supp. Fig. 5)**. Assessment of the SOD1-TdTomato signal in native-PAGE showed that dimer was only the most prominent species for WT (60%) when cells were treated with vehicle control. Other variants were predominantly monomeric, where G93A was the ALS-associated variant with the highest proportion of dimer **(Figure 4 E and F)**. Treatment with CuATSM, ebselen, or combination therapy showed increases in the proportion of dimeric SOD1 for both WT and G93A, indicating that compounds were promoting dimerization either through Cu input or disulfide formation **(Figure 4 E and F)**. Subsequent in-gel zymography showed that all variants (G85R data not shown due to no activity observed under any treatment), exhibited a significant increase in the activity of SOD1-TdTomato for CuATSM, and combination therapy treated cells compared to vehicle-treated cells **(Figure 4 E and G)**. Ebselen treatment did not appear to result in greater levels of SOD1 activity, supporting a mechanism of stabilization that is related only to disulfide formation. V148G was more enzymatically active when both CuATSM and ebselen were used in combination, as compared to when CuATSM was administered alone, perhaps due to ebselen further stabilizing the disulfide bond, which is an important PTM for SOD1 activity [64].

## Discussion

Considering that ALS-associated mutations in SOD1 disrupt its maturation, some therapeutic strategies have focused on catalysing proper SOD1 folding [30,41,65–67]. Initial methods to stabilize SOD1 were focused on promoting the formation or maintenance of the SOD1 homodimer [65,68], which was a strategy based on the success of small molecules that maintained familial amyloid polyneuropathy-associated mutants of the serum protein transthyretin in its native tetrameric conformation [69]. A caveat to the approach of promoting dimer stability for SOD1-associated ALS mutants is that SOD1 dimer formation primarily occurs when monomers are already metal replete and disulfide oxidized: a species of SOD1, which is still highly stable even when containing ALS-associated mutations [70]. Evidence points towards immature metal depleted SOD1 being a precursor to the toxic forms of misfolded or aggregated SOD1 [4,10,18,19]. Until recently, effective pharmacological chaperones targeting immature SOD1 were elusive. Ebselen is considered to have a potential duel effect on SOD1 maturation, facilitating disulfide formation, and increasing dimer affinity through binding at Cys111 [30], although it should be noted that *in vivo*, ebselen binding at Cys111 is unlikely due to the presence of reduced glutathione within cells. CuATSM is thought to facilitate the increased delivery of Cu to SOD1, increasing the pool of Cu-bound SOD1 in several SOD1 ALS animal models [31,38,40]. Here we considered that ebselen and CuATSM may be used in combination to promote proper SOD1 folding at two different points, copper binding and disulfide formation in effect acting as a CCS mimetic.

Ebselen was found to be more effective at attenuating the inclusion formation and toxicity of A4V when compared to G85R. Previous investigations have shown that SOD1 with Cu bound is less susceptible to disulfide reduction [59,71,72]. Since A4V is a WTL mutant that retains Cu-binding capacity similar to that of WT, it would be expected that Cu-bound forms of A4V would respond more favorably to ebselen-associated disulfide formation. Indeed, our data showed that combination treatment resulted in a greater level of enzymatically active V148G as compared to either CuATSM or ebselen alone, suggesting a strong effect of polytherapy to stabilise even the most destabilizing ALS-associated mutants. In contrast, the pool of G85R that binds Cu is relatively low [52,73], meaning that the synergistic effect of Cu-binding and disulfide formation that we observed for both V148G and A4V would not be expected to be seen for G85R. In line with this, we saw no additive or synergistic effect of CuATSM in combination with ebselen against G85R expression in our cell model.

Other pharmacological chaperones, such as those targeted against tryptophan-32 in SOD1 [28,29,74] or the molecular tweezer CLR01 [75], may result in more effective reduction of inclusion formation and toxicity in the case of SOD1 MBR mutants such as G85R. Likewise, derivatives of ebselen may also prove more effective at reducing inclusion formation and toxicity in the case of SOD1 MBR mutants [42]. Both ebselen and CuATSM are currently in clinical trials. CuATSM is currently in clinical trials against ALS [NCT04082832 and NCT02870634] and ebselen is in clinical trials as a potential treatment for noise-induced hearing loss [76,77], and is even being considered as a potential treatment for COVID-19 [78]. That these two compounds have available safety profiles is promising for their application as a polytherapy for ALS patients carrying SOD1 mutations. The effectiveness of CuATSM and ebselen for other forms of ALS is currently not well understood. CuATSM has been found to inhibit the paraquat-induced cytoplasmic localization of TAR DNA-binding protein 43 (TDP-43) into stress granules [79] and to prevent TDP-43 phosphorylation and fragmentation *in vivo* [80]. Ebselen is yet to be examined against TDP-43-associated forms of ALS.

Collectively, we have shown here that a biophysical understanding of the folding pathway of a protein can be exploited to target it at key points to promote proper folding. However, the strategy presented here is by no means the only one that may be pursued against SOD1-fALS. Considering the concept of proteostasis (protein homeostasis) incorporates protein synthesis, protein folding, protein trafficking, and protein degradation [81], there is potential to establish combination therapies against SOD1-fALS at multiple points. These therapies may upregulate the cellular chaperone networks [82], protein degradation pathways [83], or reduce SOD1 synthesis [84]. For example, the most clinically promising therapeutic for SOD1-associated ALS are antisense oligonucleotides (ASOs) [84], which bind to and enhance the degradation of SOD1 mRNA, effectively reducing the concentration of SOD1 within a cell. Considering SOD1 deficiency may have negative effects in animals and humans [85], complete knockdown of SOD1 via ASOs may potentially lead to complications. A different strategy may include partial knockdown of SOD1 paired with supplemental treatment with pharmacological chaperones such as CuATSM and ebselen to ensure the SOD1 that is synthesised folds properly. Indeed, cancer researchers have made marked advances in establishing polytherapies as a highly effective method by which to treat cancer [55]. Finally, future work against SOD1-associated ALS, and even other neurodegenerative diseases, should incorporate our growing knowledge of the root mechanisms and downstream effects into therapy design. We hypothesize that further improvement and expansion of the number of pharmacological chaperones that promote SOD1 folding will result in better outcomes in preclinical models and patients.

## Conflict of Interest Statement

The authors declare no conflict of interest.

## Author Contributions

Conceptualization: LM, GSAW, JJY; **Methodology:** LM, GSAW, JJY; **Validation:** LM, VS, GSAW, JJY; **Formal analysis:** LM, VS, GSAW; **Investigation:** LM, VS, GSAW; **Resources:** GSAW, JJY; **Data curation:** LM, VS, GSAW; **Writing - original draft:** LM, VS, GSAW; **Writing - review & editing:** LM, VS, GSAW, JJY; **Visualization:** LM, VS, GSAW; **Supervision:** LM, JJY; **Project administration:** LM, JJY; **Funding acquisition:** LM, GSAW, JJY.

## Acknowledgments

LM is the Bill Gole MND Fellow (Motor Neurone Disease Research Australia). GSAW is funded by the Motor Neurone Disease Association, UK, Wright/Oct18/969-799. The authors acknowledge the facilities and technical staff of the Illawarra Health and Medical Research Institute. The authors acknowledge the facilities, the technical and scientific assistance of the Fluorescence Analysis Facility in Molecular Horizons, Faculty of Science, Medicine and Health, University of Wollongong.

## Supporting Information

### Methods and Materials

#### Plasmids for Mammalian Protein Expression

Vectors for the expression of C-terminally EGFP-tagged SOD1 variants WT, A4V, C6G, G37R, H46R, D90A, G93A, E100G, G127X, and V148G on a pEGFP-N1 backbone have been previously described [3,41,47]. Plasmids for the expression of human SOD1-A4V from *Escherichia coli* have been described previously [30]. Plasmids were heat transformed into subcloning efficiency chemically competent *Escherichia coli* DH5α cells (Thermofisher, USA) and purified using miniprep kits (Thermofisher, USA) and maxiprep kits (Qiagen, Germany) as per the manufacturer’s instructions.

#### Mammalian Tissue Culture and Transfection

NSC-34 [86] cells were cultured in Dulbecco’s modified Eagle’s medium-F12 (DMEM-F12) (Invitrogen, USA), supplemented with 10% (v/v) heat inactivated fetal bovine serum (FBS) (Bovogen, Australia). In order to passage and plate NSC-34 cells, they were washed once with pre-warmed DMEM-F12 and treated with 0.25% trypsin, 0.02% EDTA dissociation reagent (Invitrogen, USA) to lift off the adherent cells. The cells were pelleted via centrifugation (300 × *g* for 5 min) and resuspended in pre-warmed culture media. Following washing, plates were seeded at a confluency of 40% and cultured at 37 °C in a humidified incubator with 5% atmospheric CO_2_ for 24 h prior to transfection (~70-80% confluent). Cells were transfected with plasmid DNA (0.5 μg per well of a 24-well plate, 2.5 μg per well of a 6-well plate) 24 h post-plating using TransIT-X2 reagent (Mirus Bioscience, USA) according to the manufacturer’s instructions.

#### Crystal Violet Assay for Cell Density

The half maximal inhibitory concentration (IC_50_) of various drugs on untransfected NSC-34 cells was determined via a crystal violet assay (0.5 g/L crystal violet, 1 % methanol (v/v), 1× PBS) as previously described [87]. NSC-34 cells were treated for 48 h with ebslen and CuATSM at concentrations ranging from 0-500 μM with a final concentration of 1% (v/v) (dimethyl sulfoxide (DMSO) (Sigma Aldrich, USA)). The drug treated NSC34 cells were then fixed via the addition of pre-warmed 4% paraformaldehyde (PFA) in 1× PBS and incubated for 30 min at room temperature prior to the addition of crystal violet solution. Crystal violet stained cells were imaged using a Gel Doc XR+ gel imager (BioRad, USA). Glacial acetic acid (100 μL 33% (v/v)) was used to release the crystal violet stain back into solution for quantification of absorbance at 590 nm on a POLARstar plate reader (BMG Labtech, Germany). The resulting data were plotted via Prism (GraphPad PRISM, Version 5.00 or Version 8.00) using a log (inhibitor) *vs.* normalized response variable slope fit.

#### Preparation of Plates for Fluorescence Microscopy

NSC-34 cells were plated into 6-well culture plates at a confluency of 40% and incubated overnight at 37 °C in a humidified incubator with 5% atmospheric CO_2_. To overcome the effect of transfection efficiency differences within assays, cells were transfected as described above and incubated for 5 h at 37 °C in a humidified incubator with 5% atmospheric CO_2_. Following incubation, cells were aspirated or lifted off with trypsin/EDTA dissociation reagent and replated into 96-well culture plates at a confluency of 30% in the presence or absence of various compounds in a final volume of 100 μL and incubated for 48 h. Following incubation, cells in culture medium were fixed via addition of 100 μL pre-warmed 4% paraformaldehyde (PFA) in 1× PBS with a 30 min incubation. Following fixation, cells were permeabilized using 0.1% Triton X-100 in 1× PBS for 5 min, which was followed by a 5 min incubation in 1× PBS with a 1:5000 dilution of Hoescht 33342 (Life Technologies, USA). Finally, cells were washed twice in 1× PBS before being immediately imaged or stored in the dark at 4 °C. Stored cells were imaged no later than 3 days after fixation.

#### Fluorescence Microscopy

A LionHeartFX automated microscope (Biotek Agilent, USA) running Cytation software (Biotek Agilent, versions 3.04 and 3.08) was used for plate-based image acquisition. NSC-34 cells expressing SOD1 mutant EGFP-tagged constructs were excited via illumination with a 465 nm LED and the emission was filtered through a 469/525 nm bandpass filter cube. All images were taken using an UPLFLN PH 10× 0.3NA objective (Olympus, Japan). Each well was imaged in a 4×4 or 5×5 tile scan. No image overlapping was used in order to avoid duplicating cell counts in later analysis stages. For imaging SOD1-WT-EGFP transfected cells, the LED power was set to 1 and integration time was 20 ms to prevent the acquisition of saturated fluorescent signal from high expressing cells. Mutant SOD1 transfected cells were imaged using an LED power of 2 and an integration time of 25 ms, set to account for variation in fluorescent intensity and lower expression. A camera gain of 10 was consistent for both WT and mutant samples. Meta data for individual wells was set to the following:

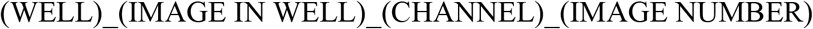

to generate a unique image identity such as B4_2_GFP_2 allowing for easy metadata assignment and data curation in image processing and analysis pipelines.

#### Image Analysis

All images generated via automated microscopy underwent pre-processing quality control to omit out of focus images and to correct illumination variation in the data sets. Out of focus images were manually assessed by users and excluded from the data set while illumination variation within images was corrected using CellProfiler software modules *Correct Illumination Calculate* and *Correct Illumination Apply*. Briefly, a 500 pixel gaussian smoothing filter was used to generate illumination functions which display the illumination variation within a set of images from one imaging session (multiple plates). The illumination function is then subtracted from the data set to correct for variation across the image. After quality control processing, images were processed in CellProfiler to segment cells within the range of 17-50 pixel units and measure intensity, granularity, size/shape, intensity distribution and texture. Accuracy of segmentation and thumbnail generation was assessed during the training of the machine learning algorithm. To determine accuracy, the user requested 100 cells of a particular “bin” and noted how many cells were incorrectly classified. This was repeated 3 times per number of cells the machine was trained on and the average accuracy noted. Once a reasonable accuracy was achieved (~97 %) all remaining cells were automatically scored via CellProfiler Analyst. Pipelines are available from the authors upon reasonable request.

#### Purified SOD1 free-thiol and homodimerization assays

Human SOD1-A4V was expressed in *Escherichia coli* BL21 (DE3) and purified as described previously [42]. The SOD1 intra-subunit disulfide bond was reduced with 40 mM dithiothreitol (DTT) overnight at 4 °C followed by desalting into N_2_ purged 20 mM Tris-HCl, 150 mM NaCl with 5 mM reduced glutathione for SEC homodimerization assays and without reduced glutathione for free-thiol assays. Compounds, including ebselen, were dissolved in DMSO and added to 20 μM SOD1-A4V at 20 or 100 μM concentration for homodimerization and free-thiol assays respectively. The reaction was incubated at 20 °C for 1 h before addition of 400 μM 4-acetamido-4’-maleimidylstilbene-2,2’-disulfonic acid and incubation at 37 °C for 90 minutes. Samples were then heated to 97 °C in non-reducing SDS sample buffer then separated by SDS-PAGE using a 15% polyacrylamide gel. Homodimerization assay samples were incubated at 20 °C for 24 h then 10 μl was loaded on an Agilent BioSEC Advance 300 Å, 4.6 × 300 mm size exclusion chromatography column along with controls for disulfide reduced and disulfide intact SOD1-A4V without ligands.

#### Immunoblotting of cell lysates

NSC-34 cells were cultured in 6-well plates, transfected, and treated with compounds similar to above methods. Importantly, following 6 h after addition of transfection complexes to cells, cells transfected with specific constructs (SOD1-WT-EGFP, SOD1-A4V-EGFP, or SOD1-G85R-EGFP) were lifted and mixed together and replated to ensure equal transfection per construct for each drug treatment. Following 48 h incubation in drugs (vehicle DMSO, 0.5 μM CuATSM, 20 μM ebselen, or a 0.5 μM CuATSM / 20 μM ebselen combination), cells were washed with prewarmed (37 °C) serum-free DMEM/F12 once, and incubated for 5 min in prewarmed 0.25% trypsin, 0.02% EDTA dissociation reagent. Once lifted, cells were harvested into microfuge tubes and pelleted at 300 × *g* for 5 min. Pellets were gently resuspended in prewarmed 1× PBS and spun again at 300 × *g* for 5 min. Supernatant was removed and cell pellets were resuspended and lysed in 100 μL ice-cold 1× Tris-buffered saline (pH 7.4) with 1% TX-100 1 mg/mL N-ethylmaleimide (NEM) supplemented with 1× Halt™ protease inhibitor (ThermoFisher, USA) to release soluble SOD1 from cells. Resuspensions were centrifuged at 20,000 × *g* for 20 min at 4 °C to pellet nuclei and insoluble material. Supernatants were carefully transferred to new microfuge tubes and these samples were flash frozen with liquid N_2_ and stored at −80 °C prior to use.

Cell lysates were defrosted on ice and mixed 1:3 with either 4× non-reducing thiol-blocking SDS-PAGE sample buffer (200 mM Tris-HCl pH 6.8, 8% SDS (w/v), 40% glycerol (v/v), 50 mM EDTA, 0.08% (w/v) bromophenol blue, 40 mM NEM) or 4× reducing SDS-PAGE sample buffer (200 mM Tris-HCl pH 6.8, 8% SDS (w/v), 40% glycerol (v/v), 50 mM EDTA, 0.08% bromophenol blue (w/v), 4% β-mercaptoethanol (v/v)). Samples for SDS-PAGE were then heated to 95 °C for 5 min prior to being loaded onto 4–20% Criterion™ TGX Stain-Free™ gels (BioRad, Australia). Gels were electrophoresed for 5 min at 100 V and then 1 h at 150 V. Following electrophoresis, total protein on the gel was quantified using a Criterion Stain Free™ Imager (BioRad, Australia) prior to transferring for immunoblotting.

Proteins separated by SDS-PAGE were transferred onto methanol activated Amersham™ Hybond™ 0.2 μm PVDF membranes (GE Healthcare, USA) at 100 V for 1 h at 4 °C using 1× transfer buffer (25 mM Tris-base 20% methanol (v/v) 192 mM glycine). Membranes were imaged with a stain-free imager post-transfer to confirm transfer and measure total protein. Following stain-free imaging, membranes were washed for 5 min in DH_2_O before being blocked in 1× Tris-buffered saline (pH 7.4) with 0.02% Tween-20 (v/v) (TBST) and 5% skim milk (w/v) at room temperature for 1 h. Following blocking, PVDF membranes were incubated in polyclonal rabbit anti-GFP primary antibody (ab290 - Abcam, USA) diluted 1:5000 in TBST with 5% skim milk (w/v) at 4 °C overnight with tipping. Following overnight incubation in primary antibody, membranes were washed three times in TBST for 10 min per wash. Following washing, membranes were incubated in goat anti-rabbit HRP-conjugated secondary antibody (P0448 - Dako, Denmark) at a dilution of 1:5000 in TBST with 5% skim milk at room temperature for 1 h with tipping. Following secondary antibody incubation, membranes were washed three times in TBST prior to chemiluminescent detection of bands using SuperSignal™ West Pico Plus substrate (Thermofisher, USA) and being imaged on a ChemiDoc™ MP Imaging System (Biorad, Australia). Analysis and quantification was performed using ImageJ (version 1.53c).

#### In-gel zymography for SOD1 enzymatic activity

Cell lysates from transfected NSC-34 cells were generated as above, with the exception that lysis buffer was 100 mM Tris-base (pH 7.5) with 0.1% TX-100 (v/v) and protease inhibitor. Cell lysates were mixed 1:2 with 3× Native-PAGE sample buffer (240 mM Tris-HCl pH 6.8, 30% glycerol (v/v), 0.03% bromophenol blue (w/v)) and loaded into Tris-glycine Native-PAGE gels (4.5% stacking gel pH 8.8, 7.5 % resolving gel pH 8.8). Samples were electrophoresed for 30 min at 60 V and then for 2.5 h at a constant voltage of 125 V at 4 °C. Following Native-PAGE, EGFP or TdTomato signal in the gel was detected using a ChemiDoc™ MP Imaging System (Biorad, Australia). Gels were then subject to zymography as described previously [63] and imaged using a GS-900™ Calibrated Densitometer (BioRad, Australia). Quantification of fluorescence signal and enzymatic activity was performed using ImageJ (version 1.53c) [88].

#### Statistical analysis

All statistical analyses are described in figure legends. All statistical analysis was performed using Prism Software version 5.00 or 8.00 (GraphPad Software, USA).

## Supporting Figures and Tables

**Supplementary Figure 1.**
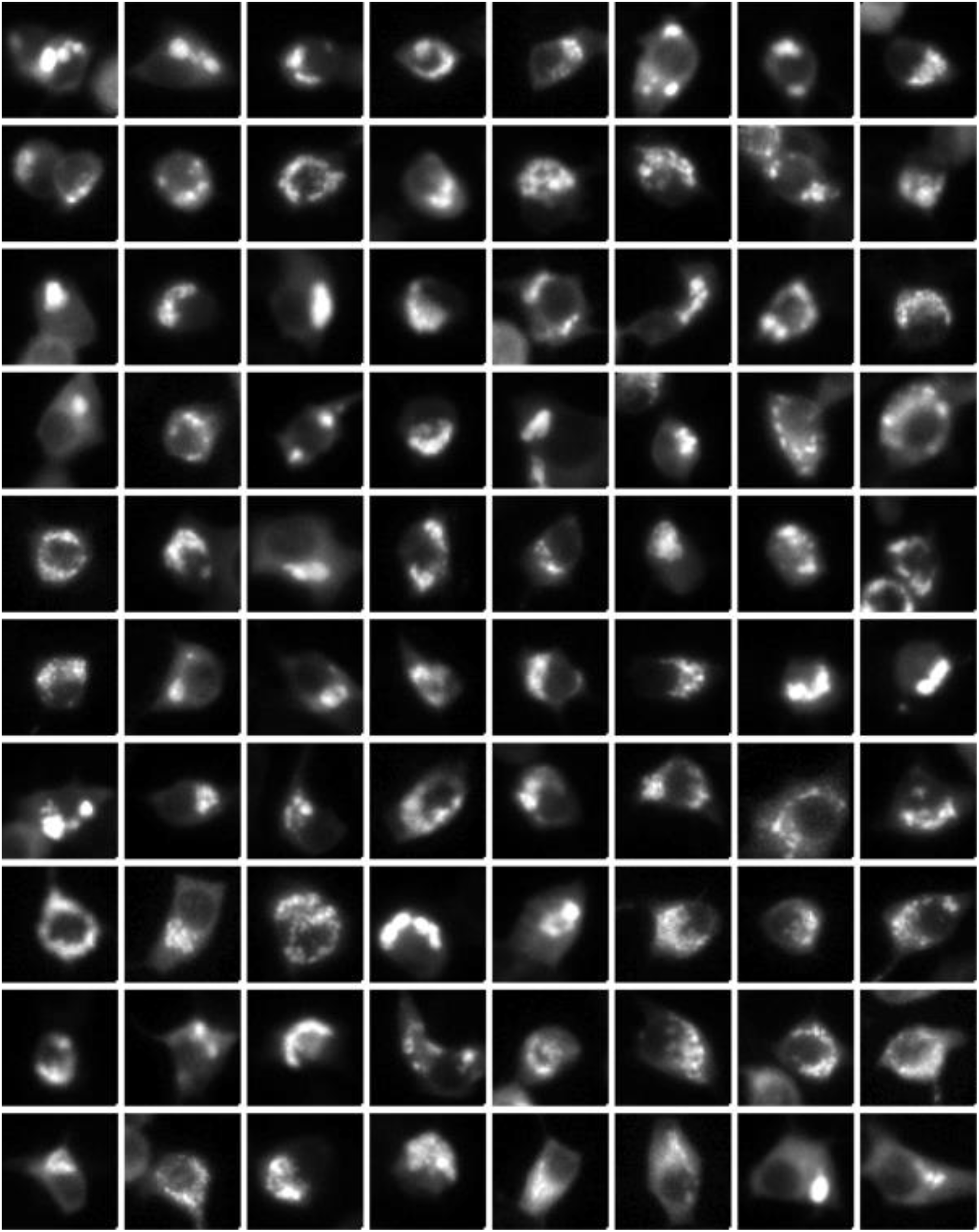
Example thumbnails of cells containing inclusions from the entire set of data used for this work, showing the heterogeneity of inclusion formation.

**Supplementary Figure 2.**
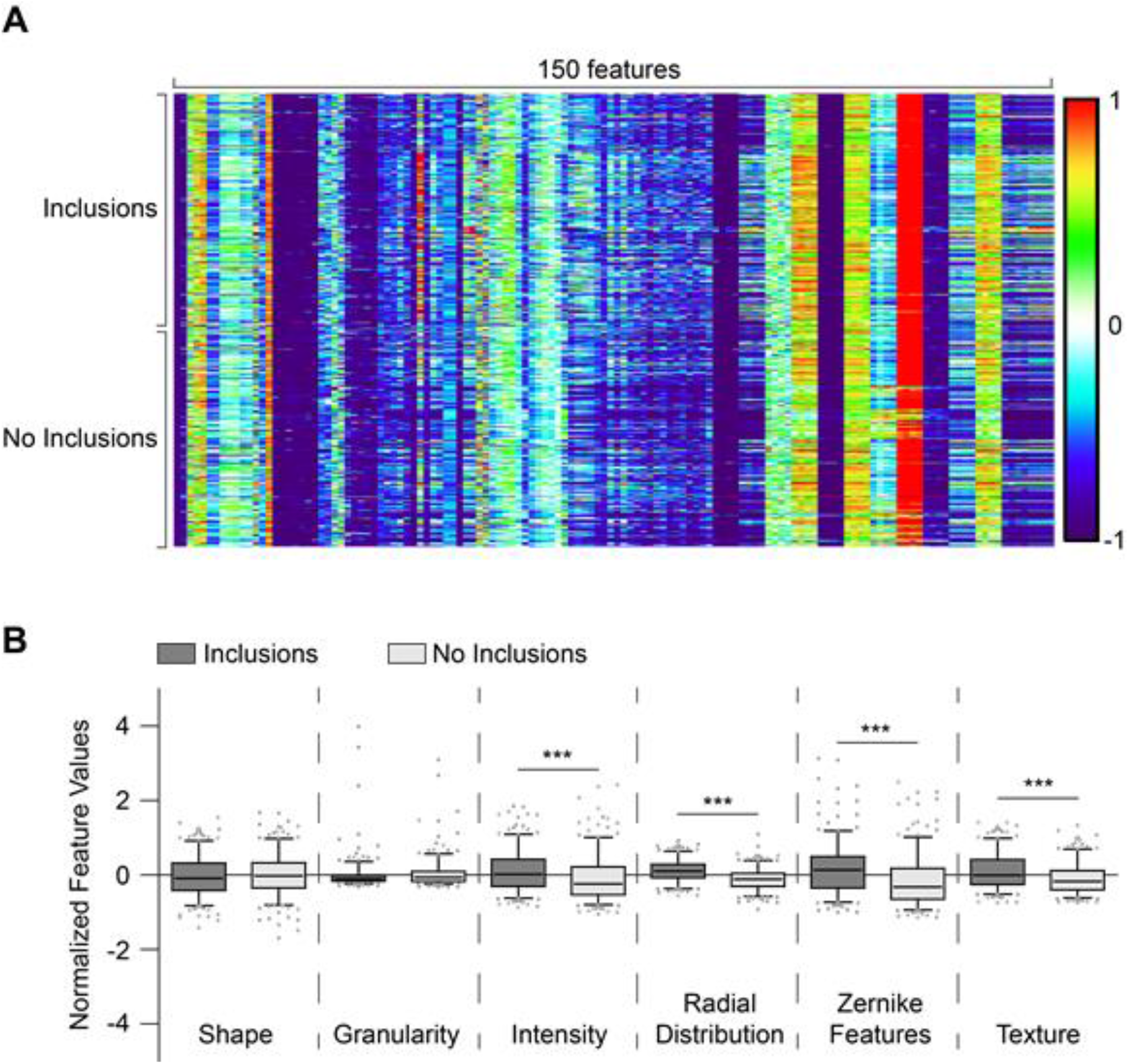
Key measurements used to generate the cytoprofiles of cells. **(A)** Heat map of the 150 features that were measured from 150 cells in each classification group (Inclusions *vs.* No Inclusions). **(B)** Z-normalised values from each extracted feature for both cells with inclusions (dark grey) and cells without inclusions (light grey). Data are plotted as box-whisker plots where whiskers show the 5-95% range, the box the 25-75% range and the middle bar the median values. Significance was determined using student’s t-test (*** *p* = *< 0.001).*

**Supplementary Figure 3.**
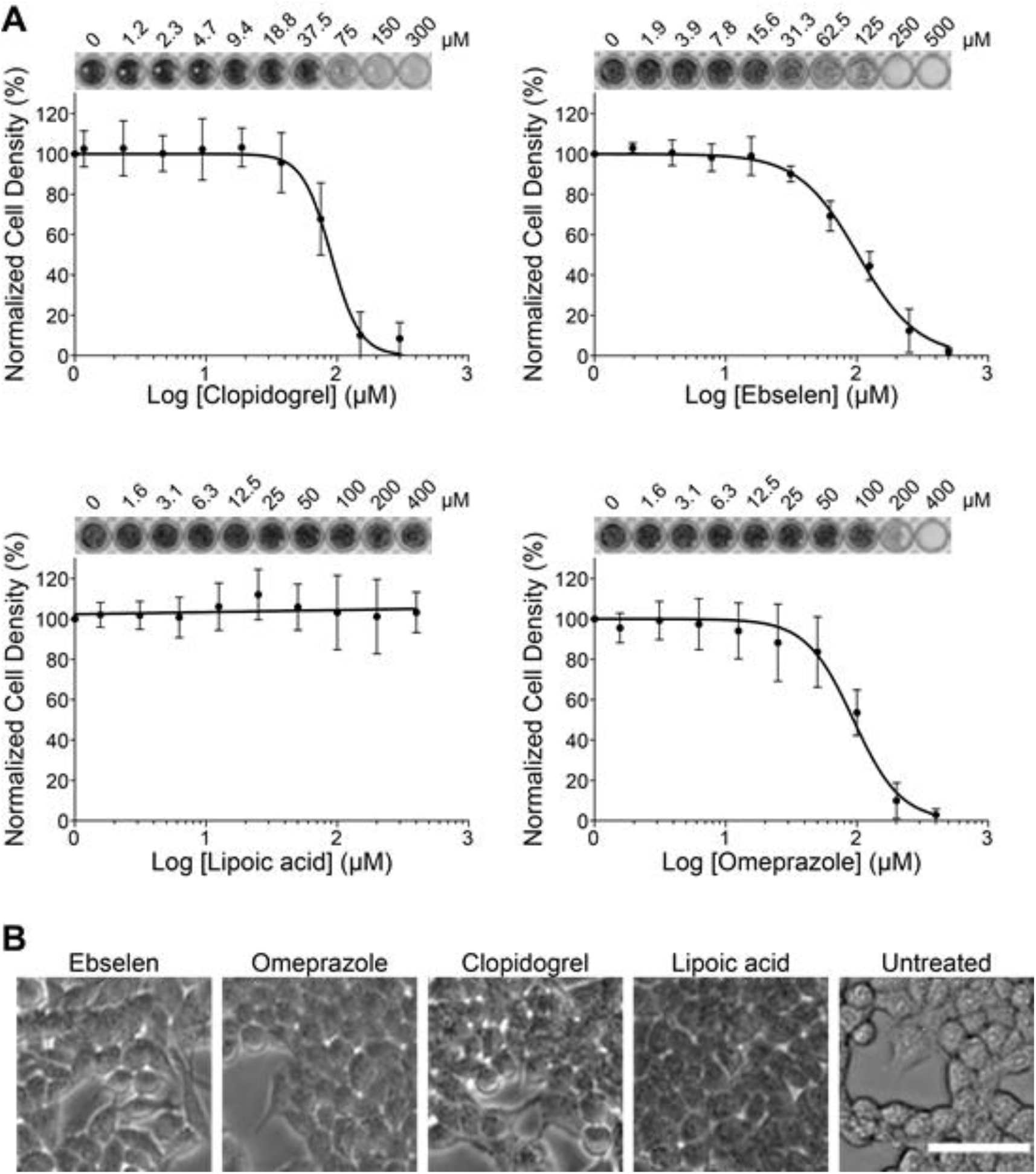
Determination of the non-toxic concentration of compounds to treat NSC-34 cells. **(A)** Crystal violet assay determined NSC-34 densities following treatment with either clopidogrel, ebselen, lipoic acid, or omeprazole. Above each plot is an example set of well images showing the crystal violet staining of cells at different compound concentrations. **(B)** Phase contrast imaging of NSC-34 cells at a concentration of 20 μ M for each drug. Scale bar = 100 μ m. Error bars represent SD of the mean from at least 3 separate biological replicates.

**Supplementary Figure 4.**
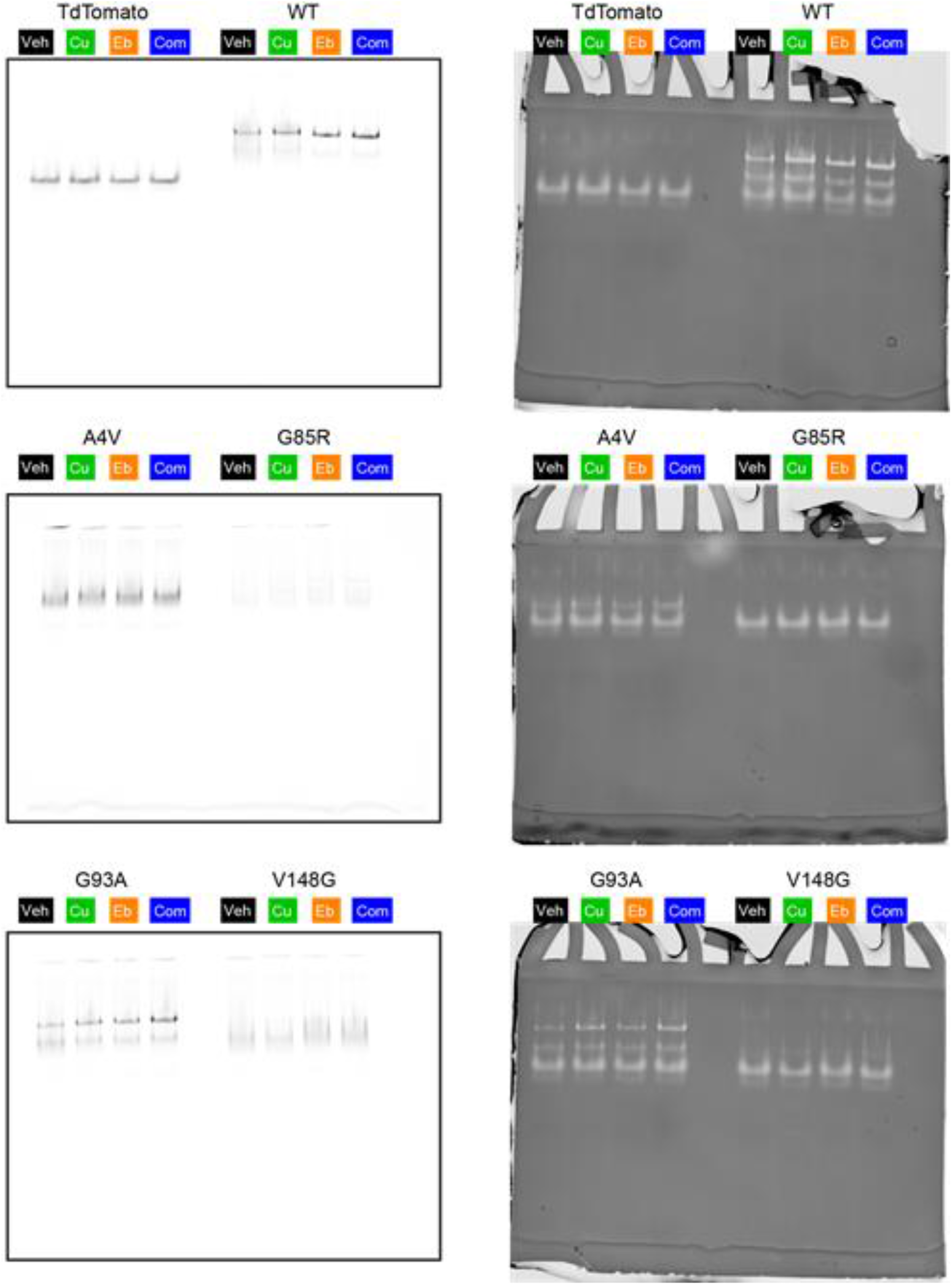
Full Native-PAGE images from TdTomato fluorescence (left) and in-gel zymography (right). Related to main text figure 6 panels E and F.

**Supplementary Figure 5.**
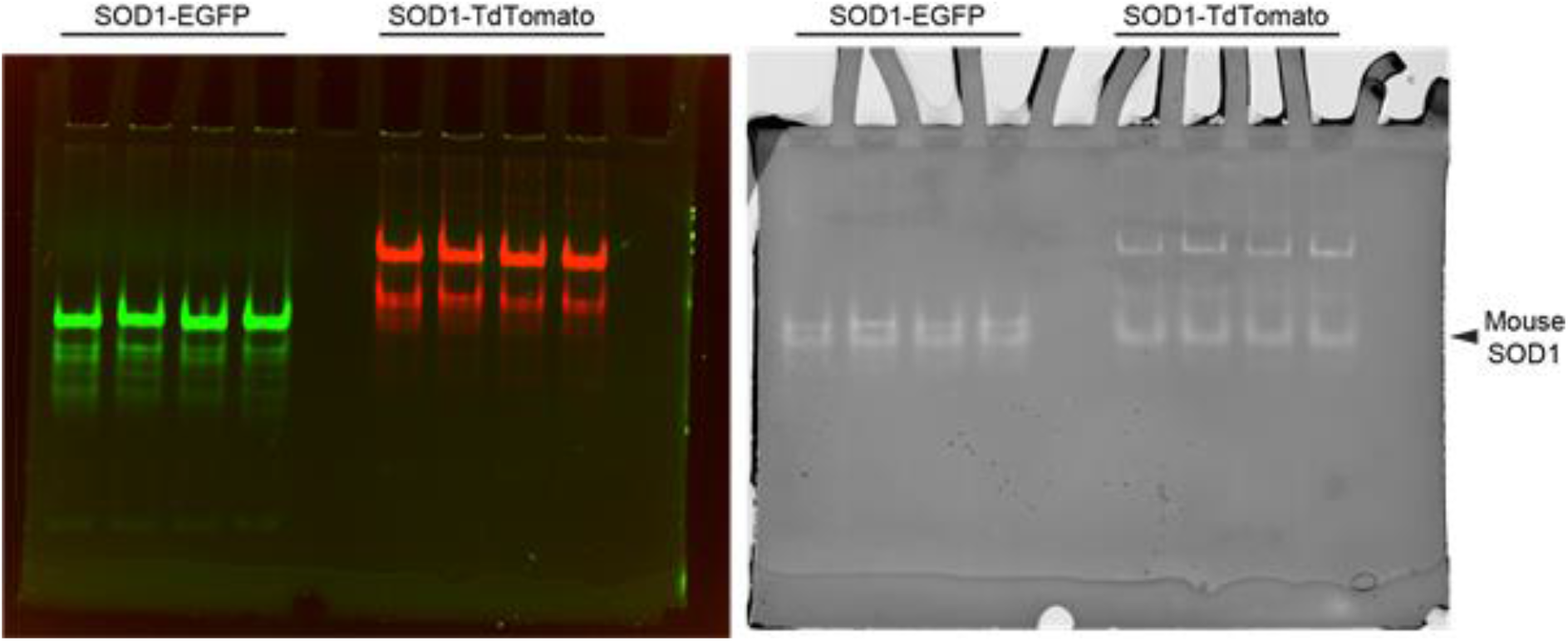
EGFP-tagged SOD1 migrates closely to mouse SOD1 in Native-gel electrophoresis. Note overexposure is to show monomer species.

**Supplementary Figure 6.**
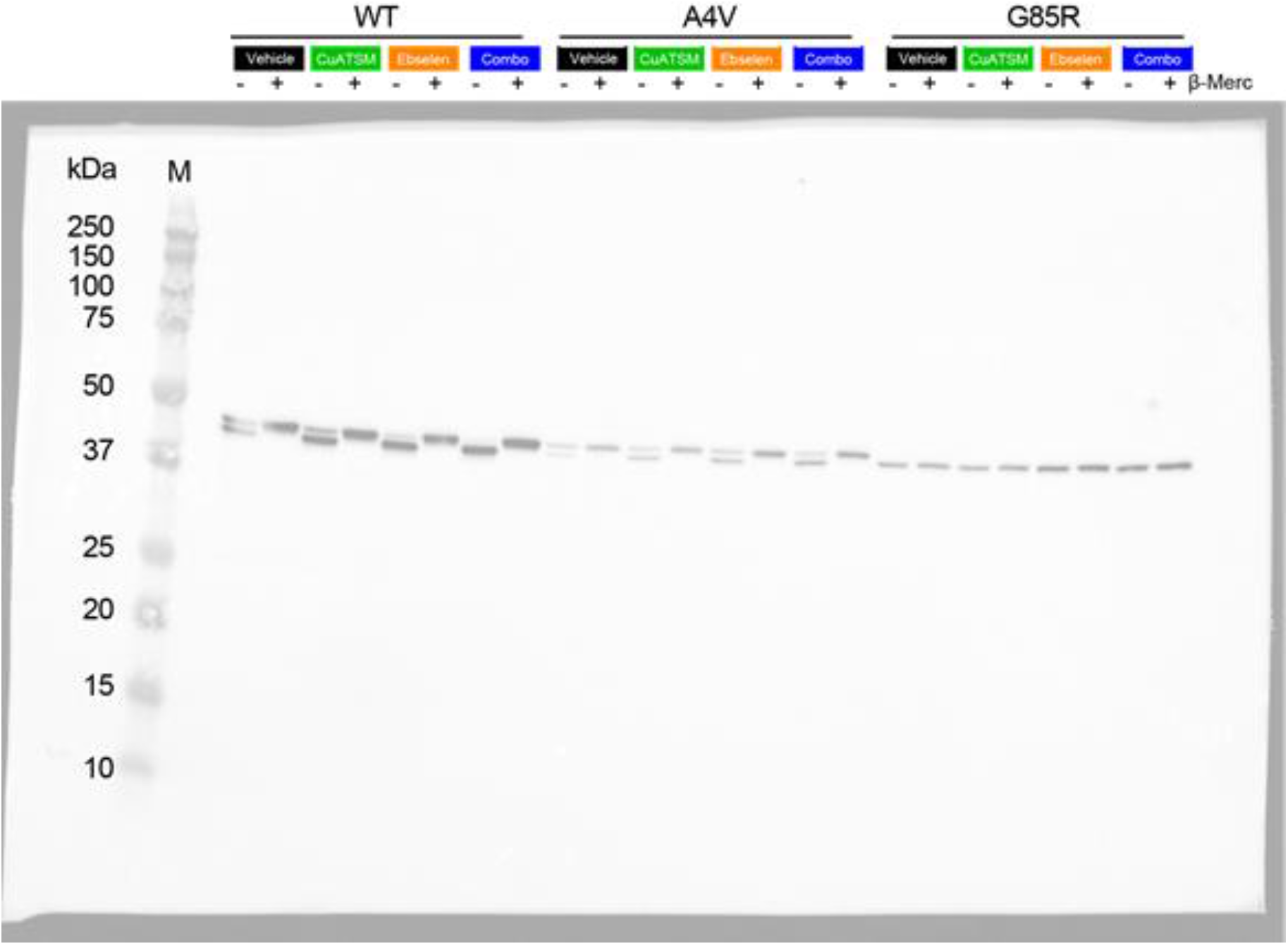
Full immunoblot representation of the electrophoretic migration assay to determine SOD1 disulfide status using SOD1-EGFP constructs. Related to main text Figure 6 panels C and D.

## Notes

### Competing Interest Statement

The authors have declared no competing interest.

